# Analysis of *DICER1* in familial and sporadic cases of Transposition of the Great Arteries

**DOI:** 10.1101/125633

**Authors:** Nelly Sabbaghian, M. Cristina Digilio, Gillian M Blue, David S. Winlaw, William D Foulkes

**Affiliations:** Lady Davis Institute, Segal Cancer Centre, Jewish General Hospital, Montreal, Quebec, Canada; Department of Medical Genetics, Bambino Gesù Pediatric Hospital, Rome, Italy; Heart Centre for Children, The Children’s Hospital at Westmead, Westmead, New South Wales 2145, and University of Sydney, Australia; Program in Cancer Genetics, Departments of Oncology and Human Genetics, McGill University, Montreal, Quebec, Canada

**Keywords:** TGA, DICER1

## Abstract

**Background:** *DICER1* plays a major role in development and in generating mature microRNAs that are important in gene expression. We screened for *DICER1* mutations in a family with DICER1 syndrome and we discovered a pathogenic mutation in a child with transposition of the great arteries (TGA). In view of a report linking *DICER1* knock-out in murine cardiomyocytes to cardiac outflow defects, we investigated the involvement of DICER1 in TGA.

**Findings:** We screened 129 germline DNA samples from children with either sporadic or familial forms of TGA for *DICER1* mutations using a Fluidigm access array, followed by next-generation sequencing. We identified 16 previously reported variants (5 synonymous, 6 intronic, and 5 missense) and 2 novel variants (1 intronic and 1 missense). We did not find any apparent pathological mutation in our cohort.

**Conclusion:** Here we report that *DICER1* mutations do not appear to play a major role in TGA.

## Findings

DICER1 is an endoribonuclease that plays an essential role in modulating the expression of genes by producing mature microRNAs (miRNA), which are small, single stranded RNA molecules that bind to and thereby inhibit target mRNAs. *DICER1*-related diseases are referred to collectively as DICER1 syndrome and result from germline mutations in individuals with rare childhood cancers such as: pleuropulmonary blastoma, cystic nephroma, Sertoli-Leydig cell tumor, embryonal rhabdomyosarcoma and other rare tumors[1] [. Several years ago, we identified a deleterious germline *DICER1* mutation (c.2117-1G>A, in intron 13 at the junction with exon 14, predicted to result in p.Gly706Aspfs*8) in a child with transposition of the great arteries (TGA), associated with a bicuspid pulmonary valve, an atrial septal defect and a patent ductus arteriosus [2]]. Later, at the age of 18, he developed a solitary nodule in the left lobe of the thyroid gland. Two years later, he was found to have further nodules and cysts in the same lobe. Other mutation-carrying persons in the family also had phenotypes consistent with the DICER1syndrome [2]. Saxena and Tabin had reported cardiac outflow defects in mice with a conditional knock-out of Dicer in the developing murine heart [3]. These two observations prompted us to screen for *DICER1* mutations in familial and sporadic cases with TGA. TGA is a cyanotic congenital heart defect (CHD) characterized by ventriculo-arterial discordance and represents 5 to 7% of CHD [4]. It is often accompanied by other structural changes that allow mixing of oxygenated and de-oxygenated blood, although there have been studies looking for the genetic causes of TGA, data so far have been inconclusive[5].

We screened 129 germline DNA samples from children with sporadic (n = 91) or familial (n = 38) forms of TGA for *DICER1* mutations using a Fluidigm access array, followed by next-generation sequencing and confirmatory Sanger sequencing [6]. Eighty-two cases were from Australia and 47 were from Italy. Details of the cases studied are shown in Supplementary Table 1. All patients signed an IRB-approved consent form.

No *DICER1* variants were detected in 110 cases. Nineteen individuals had one or more variants for a total of 5 synonymous, 7 intronic and 6 missense variants (Supplementary Table 2). c.307+13T>C and c.4886C>T are novel intronic and missense variants, respectively. c.4886C>T results in a protein with an amino acid change at position 1629, from serine to proline, (p.S1629L). SIFT (Sorting Intolerant from Tolerant)[7] and Polyphen 2 (Polymorphism Phenotyping-2)[8] predicted this variant to be “tolerated and benign”, respectively. Predictions for the other missense variants varied from possibly damaging to benign by Polyphen 2, but all variants identified were predicted to be tolerated by SIFT (supplementary table 2). No definitively damaging mutations in *DICER1* were found. In particular, we did not identify any mutations predicted to result in a truncated protein. Thus far, most disease-associated germline mutations in *DICER1* are predicted to truncate the protein [1].

This study suggests that TGA is not caused by *DICER1* mutations in humans. The full spectrum of phenotypes associated with *DICER1* mutations is still being defined, and newly-associated phenotypes such as pituitary blastoma[6] and macrocephaly [9] are still emerging. As such, it is important to fully explore all possible associations. Here we report that TGA does not appear to be part of the DICER1 syndrome. The genetics of TGA remain enigmatic [4] and it is likely that whole genome approaches in a large series of cases will be required to identify causal variants and genetic modifiers.

## List of abbreviations

TGA: Transposition of the great arteries
miRNA: microRNAs
CHD: congenital heart defect

## Declarations

### Ethics approval and consent to participate

All patients signed an IRB-approved consent form to participate in the study.

### Consent for publication

Not Applicable.

### Availability of data and material

The datasets used and/or analysed during the current study available from the corresponding author on reasonable request.

## Authors’ contributions

NS analyzed and validated the results, and wrote the manuscript. WDF wrote the manuscript with NS, and oversaw the study. MCD, GMB, and DSW provided the samples. The manuscript was reviewed and edited by all authors, who commented on and approved the final version. All authors read and approved the final manuscript.

## Acknowledgments

This work was funded by Alex’s Lemonade Stand Foundation. We thank Dr Catherine Choong, Perth, Australia, for drawing our attention to the possible connection between TGA and DICER1, and for providing updated clinical information on the patient in question. We also thank Mr François Plourde, Montreal, for his help in coordinating reception of the DNA samples, Nancy Hamel and Mona Wu, Montreal, for their help in editing the manuscript.

---

**Supplementary Table 1:** *DICER1* screening results in 47 Italian and 82 Australian germline DNA samples with clinical information.

| Case ID | Variant | Status | Clinical Info | Country of Origin |
| --- | --- | --- | --- | --- |
| 1 | negative | Sporadic | Malformation of outflow tracts | Italy |
| 2 | negative | Sporadic | Malformation of outflow tracts | Italy |
| 3 | negative | Sporadic | Malformation of outflow tracts | Italy |
| 4 | negative | Sporadic | Malformation of outflow tracts | Italy |
| 5 | negative | Sporadic | Malformation of outflow tracts | Italy |
| 6 | negative | Sporadic | Malformation of outflow tracts | Italy |
| 7 | negative | Sporadic | Malformation of outflow tracts | Italy |
| 8 | negative | Sporadic | Malformation of outflow tracts | Italy |
| 9 | c.5504A>C p.Y1835S rs747510783 | Sporadic | Malformation of outflow tracts | Italy |
| 10 | negative | Sporadic | Malformation of outflow tracts | Italy |
| 11 | negative | Sporadic | Malformation of outflow tracts | Italy |
| 12 | negative | Sporadic | Malformation of outflow tracts | Italy |
| 13 | negative | Sporadic | Malformation of outflow tracts | Italy |
| 14 | c.1935G>A p.P645P rs61751177 | Sporadic | Malformation of outflow tracts | Italy |
| 15 | c.278G>A p.G93E rs776219930 | Sporadic | Malformation of outflow tracts | Italy |
| 16 | negative | Sporadic | Malformation of outflow tracts | Italy |
| 17 | c.5364+18C>T rs777415635 | Sporadic | Malformation of outflow tracts | Italy |
| 18 | negative | Sporadic | Malformation of outflow tracts | Italy |
| 19 | negative | Sporadic | Malformation of outflow tracts | Italy |
| 20 | negative | Sporadic | Malformation of outflow tracts | Italy |
| 21 | c.1377-4T>G rs192490028 | Sporadic | Malformation of outflow tracts | Italy |
| 22 | negative | Sporadic | Malformation of outflow tracts | Italy |
| 23 | negative | Sporadic | Malformation of outflow tracts | Italy |
| 24 | negative | Sporadic | Malformation of outflow tracts | Italy |
| 25 | negative | Sporadic | Malformation of outflow tracts | Italy |
| 26 | c.2718C>T p.R906R rs370692165 | Sporadic | Malformation of outflow tracts | Italy |
| 27 | negative | Sporadic | Malformation of outflow tracts | Italy |
| 28 | negative | Sporadic | Malformation of outflow tracts | Italy |
| 29 | negative | Sporadic | Malformation of outflow tracts | Italy |
| 30 | negative | Sporadic | Malformation of outflow tracts | Italy |
| 31 | negative | Sporadic | Malformation of outflow tracts | Italy |
| 32 | negative | Sporadic | Malformation of outflow tracts | Italy |
| 33 | negative | Sporadic | Malformation of outflow tracts | Italy |
| 34 | negative | Sporadic | Malformation of outflow tracts | Italy |
| 35 | negative | Sporadic | Malformation of outflow tracts | Italy |
| 36 | negative | Sporadic | Malformation of outflow tracts | Italy |
| 37 | negative | Sporadic | Malformation of outflow tracts | Italy |
| 38 | negative | Sporadic | Malformation of outflow tracts | Italy |
| 39 | negative | Sporadic | Malformation of outflow tracts | Italy |
| 40 | negative | Sporadic | Malformation of outflow tracts | Italy |
| 41 | negative | Sporadic | Malformation of outflow tracts | Italy |
| 42 | negative | Familial | Malformation of outflow tracts* | Italy |
| 43 | c.1935G>A p.P645P rs61751177 | Familial | Ventricular Septal Defect* | Italy |
| 44 | negative | Familial | Malformation of outflow tracts# | Italy |
| 45 | negative | Familial | Tetralogy of Fallot# | Italy |
| 46 | negative | Familial | Malformation of outflow tracts^ | Italy |
| 47 | negative | Familial | Atrial Septal Defect^ | Italy |
| 48 | negative | Sporadic | Malformation of outflow tracts | Australia |
| 49 | c.5145C>T p.L1715L rs139500905 | Sporadic | Malformation of outflow tracts | Australia |
| 50 | negative | Sporadic | Malformation of outflow tracts | Australia |
| 51 | negative | Sporadic | Malformation of outflow tracts | Australia |
| 52 | negative | Sporadic | Malformation of outflow tracts | Australia |
| 53 | negative | Sporadic | Functional single ventricle | Australia |
| 54 | negative | Sporadic | Malformation of outflow tracts | Australia |
| 55 | negative | Sporadic | Malformation of outflow tracts | Australia |
| 56 | negative | Sporadic | Functional single ventricle | Australia |
| 57 | negative | Sporadic | Malformation of outflow tracts | Australia |
| 58 | c.574-5G>A rs368253792 | Sporadic | Malformation of outflow tracts | Australia |
| 59 | negative | Familial | Malformation of outflow tracts | Australia |
| 60 | negative | Familial | Functional single ventricle | Australia |
| 61 | c.1935G>A p.P645P rs61751177 | Sporadic | Malformation of outflow tracts | Australia |
| 62 | c.179C>T p.T60I rs587778228 | Familial | Malformation of outflow tracts | Australia |
| 63 | negative | Familial | Functional single ventricle | Australia |
| 64 | negative | Sporadic | Functional single ventricle | Australia |
| 65 | c.1377-4T>G rs192490028 | Sporadic | Heterotaxy | Australia |
| 66 | negative | Familial | Functional single ventricle | Australia |
| 67 | negative | Sporadic | Functional single ventricle | Australia |
| 68 | negative | Sporadic | Functional single ventricle | Australia |
| 69 | negative | Familial | Malformation of outflow tracts | Australia |
| 70 | negative | Sporadic | Malformation of outflow tracts | Australia |
| 71 | negative | Sporadic | Malformation of outflow tracts | Australia |
| 72 | negative | Familial | Malformation of outflow tracts | Australia |
| 73 | negative | Sporadic | Malformation of outflow tracts | Australia |
| 74 | negative | Familial | Malformation of outflow tracts | Australia |
| 75 | negative | Familial | Malformation of outflow tracts | Australia |
| 76 | negative | Familial | Malformation of outflow tracts | Australia |
| 77 | negative | Familial | Malformation of outflow tracts | Australia |
| 78 | negative | Sporadic | Malformation of outflow tracts | Australia |
| 79 | negative | Familial | Malformation of outflow tracts | Australia |
| 80 | negative | Familial | Malformation of outflow tracts | Australia |
| 81 | negative | Sporadic | Malformation of outflow tracts | Australia |
| 82 | c.2337A>G p.T779T rs747210633 | Familial | Malformation of outflow tracts | Australia |
| 83 | c.2040+29T>C rs370866625 | Familial | Malformation of outflow tracts | Australia |
|  | c.3458G>A p.C1153Y rs762999390 |  |  |  |
|  | c.5527+19A>G rs765497219 |  |  |  |
| 84 | negative | Familial | Heterotaxy | Australia |
| 85 | negative | Familial | Malformation of outflow tracts | Australia |
| 86 | negative | Sporadic | Malformation of outflow tracts | Australia |
| 87 | negative | Sporadic | Malformation of outflow tracts | Australia |
| 88 | negative | Familial | Functional single ventricle | Australia |
| 89 | negative | Sporadic | Malformation of outflow tracts | Australia |
| 90 | negative | Sporadic | Malformation of outflow tracts | Australia |
| 91 | c.3093+149_3093+153delGTTTT<br>rs575610432 | Sporadic | Malformation of outflow tracts | Australia |
| 92 | negative | Familial | Malformation of outflow tracts | Australia |
| 93 | negative | Sporadic | Functional single ventricle | Australia |
| 94 | negative | Sporadic | Malformation of outflow tracts | Australia |
| 95 | c.1935G>A p.P645P rs61751177 | Sporadic | Malformation of outflow tracts | Australia |
| 96 | negative | Sporadic | Malformation of outflow tracts | Australia |
| 97 | negative | Familial | Malformation of outflow tracts | Australia |
| 98 | negative | Sporadic | Malformation of outflow tracts | Australia |
| 99 | negative | Sporadic | Malformation of outflow tracts | Australia |
| 100 | negative | Familial | Functional single ventricle | Australia |
| 101 | negative | Sporadic | Malformation of outflow tracts | Australia |
| 102 | negative | Sporadic | Malformation of outflow tracts | Australia |
| 103 | negative | Familial | Malformation of outflow tracts | Australia |
| 104 | negative | Sporadic | Malformation of outflow tracts | Australia |
| 105 | negative | Familial | Functional single ventricle | Australia |
| 106 | negative | Sporadic | Malformation of outflow tracts | Australia |
| 107 | negative | Sporadic | Malformation of outflow tracts | Australia |
| 108 | negative | Familial | Malformation of outflow tracts | Australia |
| 109 | negative | Sporadic | Malformation of outflow tracts | Australia |
| 110 | negative | Familial | Malformation of outflow tracts | Australia |
| 111 | negative | Familial | Heterotaxy | Australia |
| 112 | c.4796G>A p.R1599Q rs569615549 | Sporadic | Malformation of outflow tracts | Australia |
| 113 | negative | Familial | Heterotaxy | Australia |
| 114 | negative | Sporadic | Malformation of outflow tracts | Australia |
| 115 | c.307+13T>C | Sporadic | Malformation of outflow tracts | Australia |
|  | c.1278A>G p.E426E rs878855242 |  |  |  |
| 116 | negative | Sporadic | Malformation of outflow tracts | Australia |
| 117 | negative | Sporadic | Malformation of outflow tracts | Australia |
| 118 | negative | Sporadic | Malformation of outflow tracts | Australia |
| 119 | negative | Familial | Malformation of outflow tracts | Australia |
| 120 | negative | Sporadic | Malformation of outflow tracts | Australia |
| 121 | negative | Sporadic | Malformation of outflow tracts | Australia |
| 122 | c.4886C>T p.S1629L | Familial | Malformation of outflow tracts | Australia |
| 123 | negative | Sporadic | Malformation of outflow tracts | Australia |
| 124 | negative | Familial | Malformation of outflow tracts | Australia |
| 125 | negative | Familial | Malformation of outflow tracts | Australia |
| 126 | negative | Sporadic | Malformation of outflow tracts | Australia |
| 127 | negative | Sporadic | Malformation of outflow tracts | Australia |
| 128 | negative | Familial | Malformation of outflow tracts | Australia |
| 129 | negative | Sporadic | Malformation of outflow tracts | Australia |

\*, # and ^ are 3 pairs of cousins among the familial Italian cohort families

**Supplementary Table 2:**
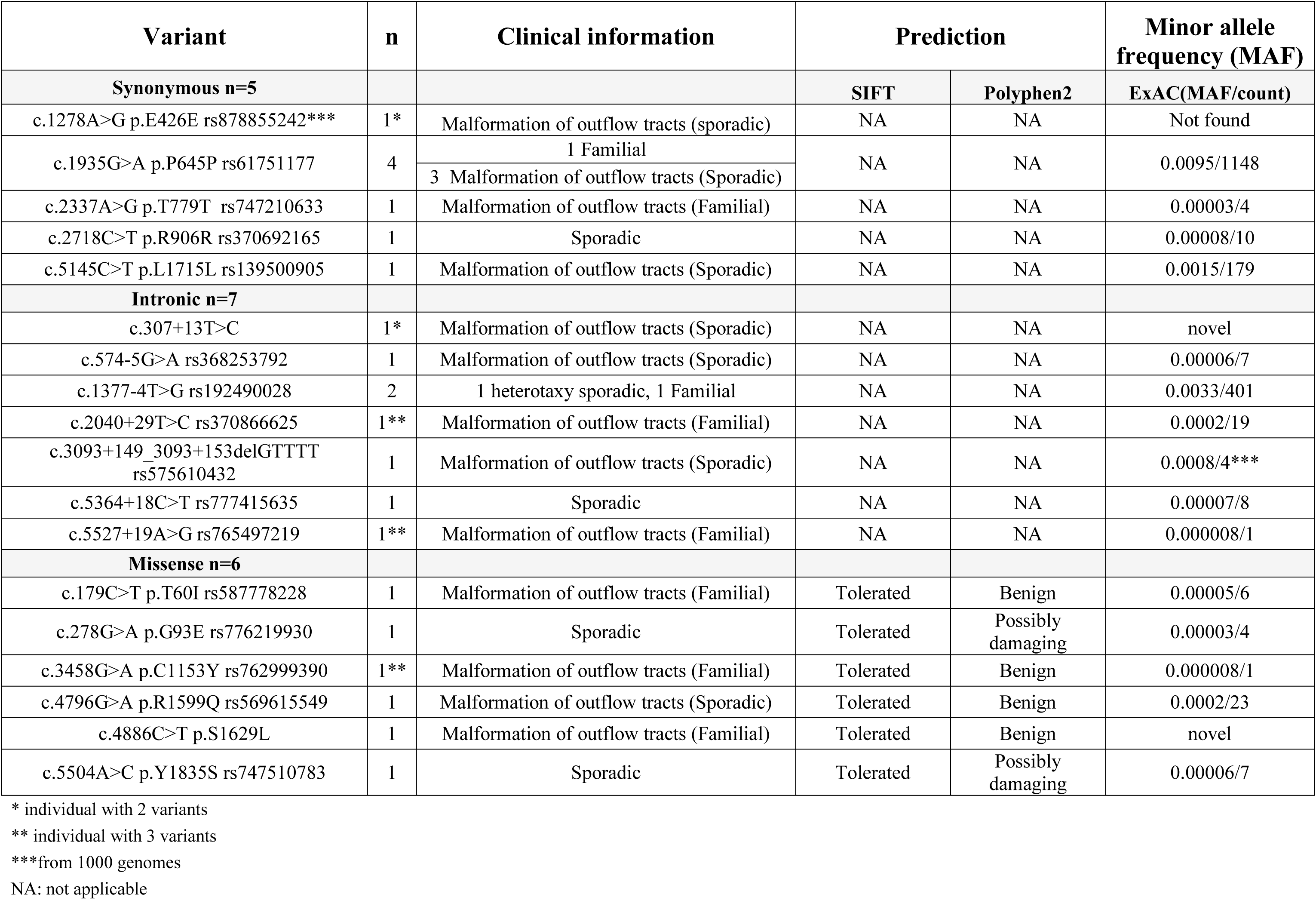
Summary of *DICER1* Variants and Predictions.

| Variant | n | Clinical information | Prediction |  | Minor allele frequency (MAF) |
| --- | --- | --- | --- | --- | --- |
| Synonymous n=5 |  |  | SIFT | Polyphen2 | ExAC(MAF/count) |
| c.1278A>G p.E426E rs878855242*** | 1* | Malformation of outflow tracts (sporadic) | NA | NA | Not found |
| c.1935G>A p.P645P rs61751177 | 4 | 1 Familial | NA | NA | 0.0095/1148 |
|  |  | 3 Malformation of outflow tracts (Sporadic) |  |  |  |
| c.2337A>G p.T779T rs747210633 | 1 | Malformation of outflow tracts (Familial) | NA | NA | 0.00003/4 |
| c.2718C>T p.R906R rs370692165 | 1 | Sporadic | NA | NA | 0.00008/10 |
| c.5145C>T p.L1715L rs139500905 | 1 | Malformation of outflow tracts (Sporadic) | NA | NA | 0.0015/179 |
| Intronic n=7 |  |  |  |  |  |
| c.307+13T>C | 1* | Malformation of outflow tracts (Sporadic) | NA | NA | novel |
| c.574-5G>A rs368253792 | 1 | Malformation of outflow tracts (Sporadic) | NA | NA | 0.00006/7 |
| c.1377-4T>G rs192490028 | 2 | 1 heterotaxy sporadic, 1 Familial | NA | NA | 0.0033/401 |
| c.2040+29T>C rs370866625 | 1** | Malformation of outflow tracts (Familial) | NA | NA | 0.0002/19 |
| c.3093+149_3093+153delGTTTT rs575610432 | 1 | Malformation of outflow tracts (Sporadic) | NA | NA | 0.0008/4*** |
| c.5364+18C>T rs777415635 | 1 | Sporadic | NA | NA | 0.00007/8 |
| c.5527+19A>G rs765497219 | 1** | Malformation of outflow tracts (Familial) | NA | NA | 0.000008/1 |
| Missense n=6 |  |  |  |  |  |
| c.179C>T p.T60I rs587778228 | 1 | Malformation of outflow tracts (Familial) | Tolerated | Benign | 0.00005/6 |
| c.278G>A p.G93E rs776219930 | 1 | Sporadic | Tolerated | Possibly damaging | 0.00003/4 |
| c.3458G>A p.C1153Y rs762999390 | 1** | Malformation of outflow tracts (Familial) | Tolerated | Benign | 0.000008/1 |
| c.4796G>A p.R1599Q rs569615549 | 1 | Malformation of outflow tracts (Sporadic) | Tolerated | Benign | 0.0002/23 |
| c.4886C>T p.S1629L | 1 | Malformation of outflow tracts (Familial) | Tolerated | Benign | novel |
| c.5504A>C p.Y1835S rs747510783 | 1 | Sporadic | Tolerated | Possibly damaging | 0.00006/7 |

\* individual with 2 variants
\*\* individual with 3 variants
\*\*\*from 1000 genomes
NA: not applicable

## References

1. Foulkes, W.D., J.R. Priest, and T.F. Duchaine, DICER1: mutations, microRNAs and mechanisms. Nat Rev Cancer, 2014. 14(10): p. 662–72.

2. Foulkes, W.D., et al., Extending the phenotypes associated with DICER1 mutations. Hum Mutat, 2011. 32(12): p. 1381–4.

3. Saxena, A. and C.J. Tabin, miRNA-processing enzyme Dicer is necessary for cardiac outflow tract alignment and chamber septation. Proc Natl Acad Sci U S A, 2010. 107(1): p. 87–91.

4. De Luca, A., et al., Familial transposition of the great arteries caused by multiple mutations in laterality genes. Heart, 2010. 96(9): p. 673–7.

5. Muncke, N., et al., Missense mutations and gene interruption in PROSIT240, a novel TRAP240-like gene, in patients with congenital heart defect (transposition of the great arteries).Circulation, 2003. 108(23): p. 2843–50.

6. de Kock, L., et al., Pituitary blastoma: a pathognomonic feature of germ-line DICER1 mutations. Acta Neuropathol, 2014. 128(1): p. 111–22.

7. Ng, P.C. and S. Henikoff, SIFT: Predicting amino acid changes that affect protein function. Nucleic Acids Res, 2003. 31(13): p. 3812–4.

8. Sunyaev, S.R., et al., PSIC: profile extraction from sequence alignments with position-specific counts of independent observations. Protein Eng, 1999. 12(5): p. 387–94.

9. Khan, N.E., et al., Macrocephaly associated with the DICER1 syndrome. Genet Med, 2016.

